# Stable Equilibria under Random Fitness and Recombination Rates

**DOI:** 10.1101/2023.11.10.566601

**Authors:** Noah Babel

## Abstract

We develop and analyze a two-locus biallelic genetic model with randomized recombination rates and fitness matrices. While deterministic models reveal a trajectory toward the simplex center before stabilizing at polymorphisms, introducing random recombination pinpointed regions of attraction dependent on the fitness matrix and recombination fraction range. As recombination randomness intensifies, transitions in these attraction regions are observed. Analysis with random fitness matrices highlighted genotype attraction towards the tetrahedron vertices with edges guiding genotypic clustering.

## 1 Introduction

The interaction between genetic linkage and selection has been a focus of evolutionary genetics since [1] first outlined the progression towards tighter linkage between two loci. The first attempt at specific analysis of the model was by [2] using a symmetric viability model. Further studies delved into the influence of natural selection and viability on the pattern of polymorphisms and linkage dynamics (Kimura, 1965; Lewontin and Kojima, 1960, respectively). Notably, much of this early work revolved around deterministic fitness frameworks with fixed recombination rates, primarily focusing on equilibrium states and their stability. While these deterministic models have provided invaluable insights, real-world genetic processes often have inherent randomness. Introducing these random dynamics, however, brings about significant complexity, posing challenges in both formulation and interpretation.

In this light, our research investigates dynamic behavior introduced by a genotypic fitness matrix and recombination fraction that undergo randomization every generational step. Employing primarily Monte Carlo simulations and probabilistic analysis, our investigation aims to understand the impact of such randomization on the dynamics of two linked biallelic gene loci.

## 2 Random two-locus symmetric viability model

Consider the standard two-locus symmetric viability model as outlined in [3]. There are two loci with two alleles each: *A* and *a* for the first locus, and *B* and *b* for the second. Let the gametic frequencies be represented as *x*_1_ for *AB, x*_2_ for *Ab, x*_3_ for *aB*, and *x*_4_ for *ab*, where 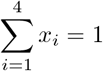. Let *w*_*ij*_ ≤ 0 denote the viability of an individual with genotype *ij*, these fitnesses are expressed in terms of a symmetric matrix

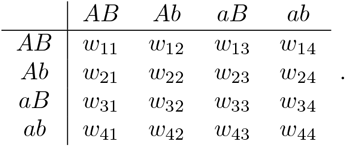

Assuming the absence of sex and position effects, we can rewrite:

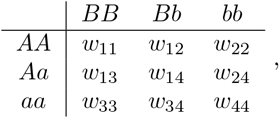

where each *w*_*ij*_ is independently and identically sampled from a uniform distribution over the interval [0, 1]. To account for the variability in meiotic recombination rates, the stochastic recombination fraction *R* is sampled at each generation from a uniform distribution over an interval within the range [0, 0.5].

The marginal mean fitnesses, represented by *W*_*i*_ can be expressed as:

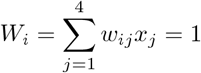

for *i* ∈{1, 2, 3, 4} . The transformation equations dictating the change in gametic frequencies from one generation to the next are:

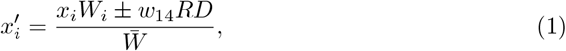

where the linkage disequilibrium function *D* = *x*_1_*x*_4_ − *x*_2_*x*_3_, mean fitness 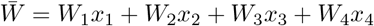, and the choice of the *±* operator for *x*_1_ and *x*_4_ is negative, while for *x*_2_ and *x*_3_ is positive.

For ease of presentation, for any given index *i* ∈ 1, 2, 3, 4, let *π* := 5 − *i* represent its corresponding ‘twin’ index, and let *r, s* = 1, 2, 3, 4 *\ i, π* denote the set of non-twin indices. This notation implies that *r* and *s* take values in 2, 3 when *i* ∈ 1, 4, and in 1, 4 when *i*∈ 2, 3. Importantly, this establishes that *w*_*iπ*_ = *w*_*rs*_ remains constant across all values of *i*. Utilizing this concise notation, we can derive the following compact representation:

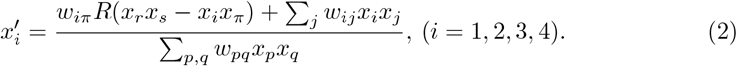

Using the above representation and the inequality

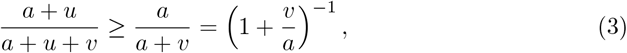

one deduces the following lower bound for the updated frequencies

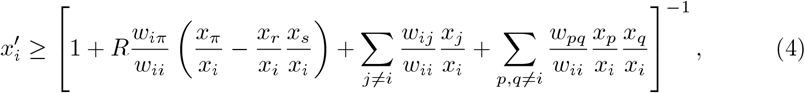

this follows by taking in (3)

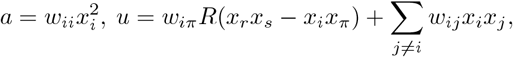

respectively

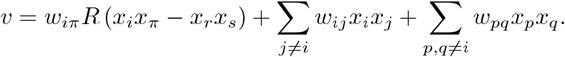

Inequality (4) plays a key role in studying the dynamics of the frequency process. It shows that, if one component *x*_*i*_ is very large compared to the other ones then a lower bound on 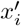 can be established. Even more, assume that 1 − *x*_*i*_ = *O*(*ε*). Then inequality (4) ensures that 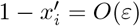, as well, which translates to *if x*_*i*_ *is close to* 1 *then* 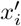 *remains close to* 1. To see that, assume for instance 1 − *x*_*i*_ ≤ *ε*, hence, for *j≠i, x*_*j*_ ≤ *ε* and

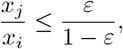

from which, one infers, based on (4), that

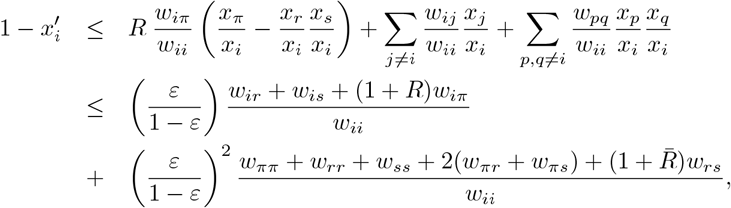

where, for convenience, we let 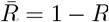. This shows that 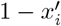 remains within the same order of magnitude as 1− *x*_*i*_.

We utilize the tetrahedral representation to depict potential chromosome frequency states using barycentric coordinates. This representation provides a geometric interpretation of the model where the distances of a point from the tetrahedron’s vertices to the opposite face are directly proportional to the gamete frequencies of that point.

## 3 Convergence behavior under fixed recombination

In deterministic scenarios, given any initial state and fixed recombination fraction, the system almost always (see Hastings 1984) deterministically converges to an equilibrium. In the case of symmetric viabilities, trajectories toward equilibrium first approach a curve defined by the pairs vertices *AB* and *ab* or *Ab* and *aB* and their polymorphisms **x**^∗^. Subsequent evolution along this curve leads the system trajectories to converge to **x**^∗^ (See Figures 1 and 2).

**Fig 1.**
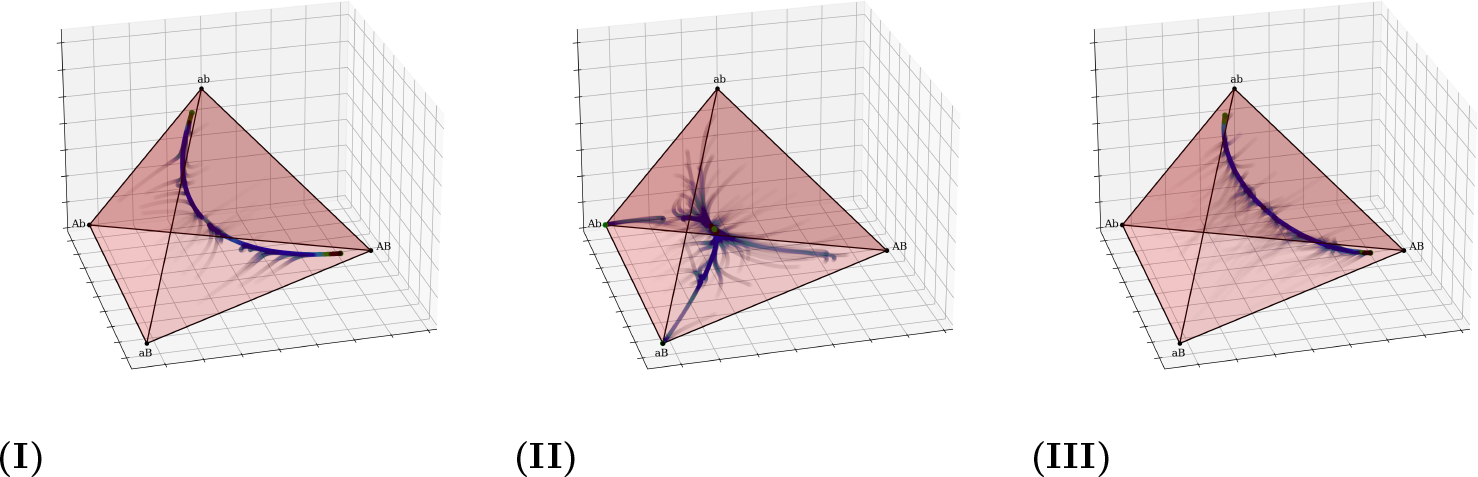
Convergent Trajectories for Numerical Simulations with Deterministic *R* The trajectory density is visualized through a heatmap with equilibrium points highlighted in green. For these and subsequent figures: roman numerals correspond to parameters in Table 1, initialization involves 10000 uniformly distributed **x**_0_, and deterministic simulations are terminated when the difference across 100 consecutive generations is less than 10^−10^.

**Fig 2.**
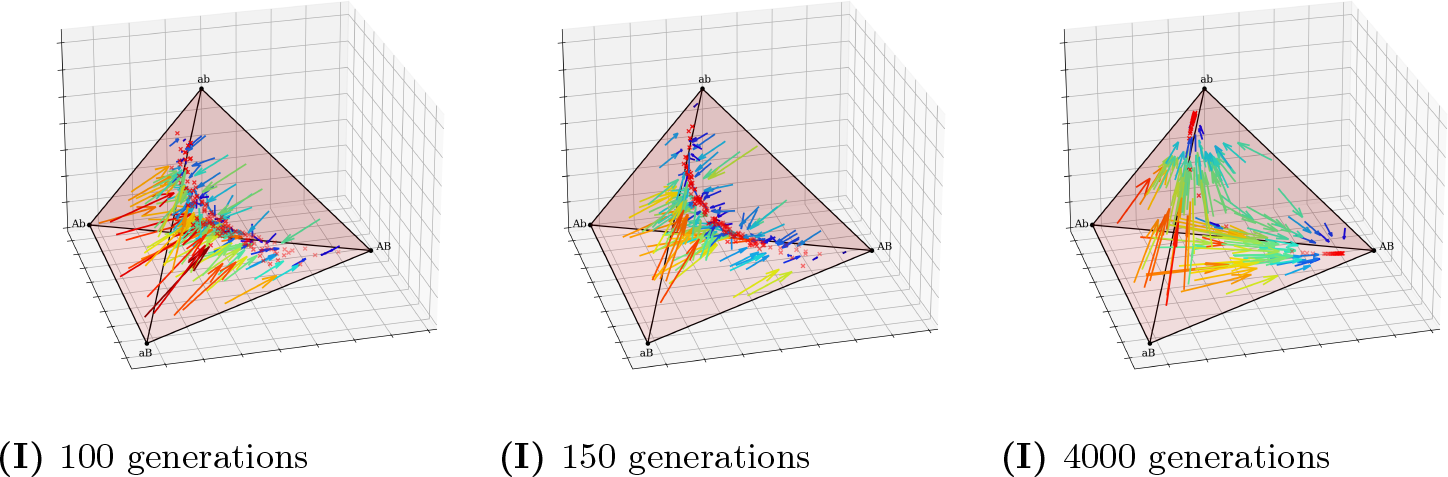
Vector Field Evolution of Trajectories For this and all subsequent vector field diagrams, the strength of a vector from a given blue sphere x within the parameter space is represented by the length and color of the corresponding vector arrow, with the region of attraction after the indicated number of generations marked by a red × symbol.

**Table 1.** Numerical Cases of Polymorphic Equilibria with Symmetric Viabilities and Deterministic *R*.

|  | Parameters | Equilibria |
| --- | --- | --- |
| I | $w_{12} = w_{34} = w_{24} = w_{13} = 0.996$<br>$w_{11} = w_{44} = 0.995$<br>$w_{22} = w_{33} = 0.97$<br>$w_{14} = 1$<br>$r = 0.05$ | 1. $\mathbf{x}^* = [0.8878, 0.0542, 0.0542, 0.0038]$<br>2. $\mathbf{x}^* = [0.0038, 0.0542, 0.0542, 0.8878]$ |
| II | $w_{12} = w_{34} = w_{24} = w_{13} = 0.995$<br>$w_{11} = w_{44} = 0.99$<br>$w_{22} = w_{33} = 0.9$<br>$w_{14} = 1$<br>$r = 0.02$ | 1. $\mathbf{x}^* = [0.866, 0.0625, 0.0625, 0.009]$<br>2. $\mathbf{x}^* = [0.009, 0.0625, 0.0625, 0.866]$ |
| III | $w_{12} = w_{34} = w_{24} = w_{13} = 0.96$<br>$w_{11} = w_{44} = 0.97$<br>$w_{22} = w_{33} = 0.98$<br>$w_{14} = 1$<br>$r = 0.04$ | 1. $\mathbf{x}^* = [0.1539, 0.6562, 0.036, 0.1539]$<br>2. $\mathbf{x}^* = [0.1539, 0.036, 0.6562, 0.1539]$<br>3. $\mathbf{x}^* = [0.8888, 0.05, 0.05, 0.0112]$<br>4. $\mathbf{x}^* = [0.0112, 0.05, 0.05, 0.8888]$ |

<sup>1</sup> Numerically determined cases of polymorphic equilibria previously derived by [3].

The trajectories approached a curved symmetric invariant set, representing a transient phase within the state space, after which trajectories then progress towards their respective equilibrium points. Indeed, the vector field for the deterministic case first directs trajectories toward the center of the tetrahedron and then directs them toward their equilibrium points (Figure 2).

## 4 Convergence behavior under random recombination

When the recombination fraction, *R* changes randomly at each generation, the trajectories are stochastic. Given a specific range of uniformly distributed values of *R*, we find regions of attraction *S* such that trajectories from all ***x***_**0**_ in *S* remain restricted with high probability.

The multicolored regions in Figure 3 denote the basin boundaries. Within these boundaries, the final equilibrium state appears to be determined, not by **x**_**0**_, but by the randomly selected *R* in the initial few generations.

**Fig 3.**
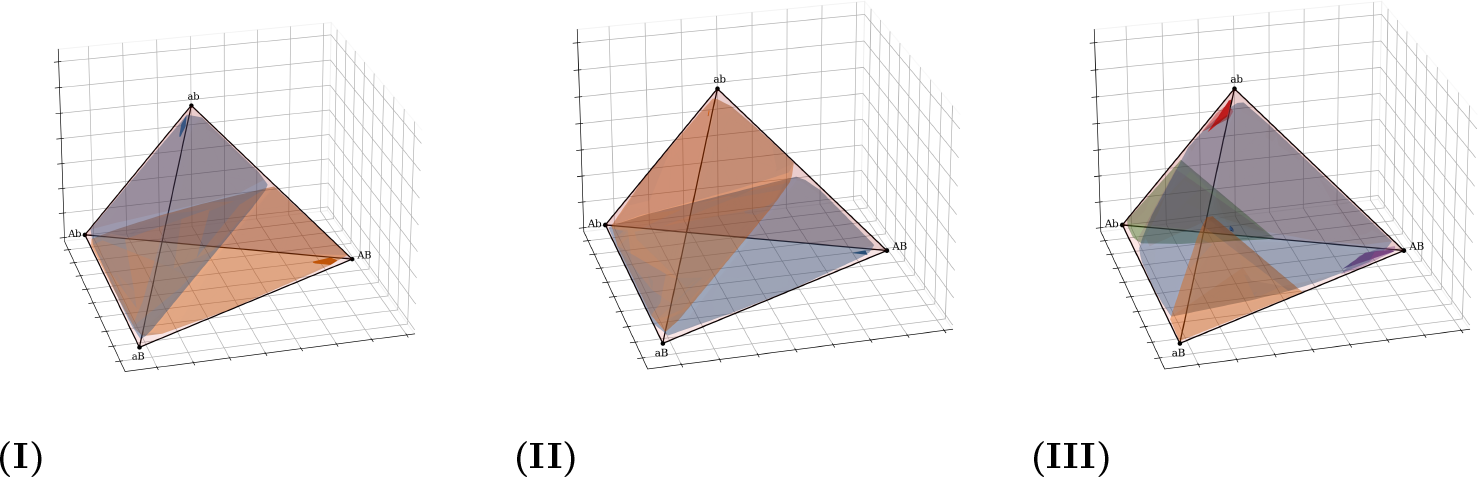
Regions of Attraction for Numerical Cases with Random *R* For this and all following random *R* diagrams: the model runs until the change across 100 consecutive generations is ⪅ 10^−4^, depending on the range of the recombination rate, *R* is uniformly chosen from a range such that its mean is the corresponding deterministic R from Table 1, and for all **x**_0_ in a given transparent domain, *S* exists as an opaque convex hull of the same color.

In this system, genotypes, while initially in a basin of attraction, can be perturbed out of it when there’s a consistent trend in the sampled *R* values over multiple generations. In the limit, such perturbations will consistently occur. Thus, when a system exhibits multiple domains of attraction within the tetrahedron’s interior, it’s more accurate to state that sequences do not truly converge to a region and instead fluctuate around a distribution. Conversely, for systems where genotype frequencies approach fixation, the frequencies of the other genotypes become so infinitesimal that in any finite population, they would be lost.

Moreover, setting progressively stricter convergence criteria by reducing the change threshold tends to prioritize regions with denser distributions. These regions exhibit slower divergence and are therefore more likely to meet stringent change thresholds. Conversely, regions that, upon stabilization, exhibit consistent albeit lower concentration of points risk being overlooked. Thus a more accurate representation of the system’s long-term behavior is to evaluate the distribution after an extensive number of generations.

In our analysis, we considered three genotypic fitness sets (see Figure 4): double heterozygote fittest, single heterozygote fittest, and double homozygote fittest, and evaluated the effects of deterministic versus random recombination fraction for each case.

**Fig 4.**
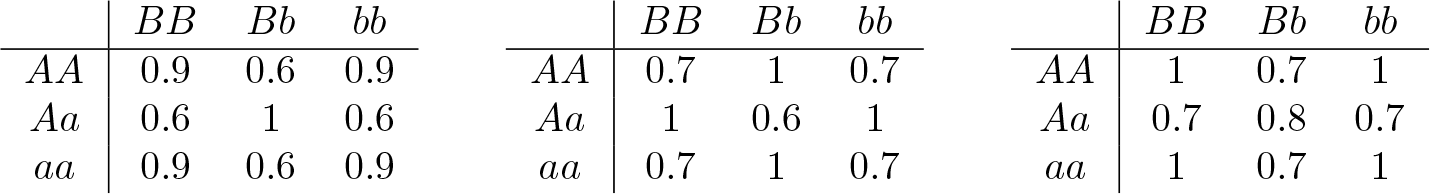
Sampled Fitness Matrices (i) Double Heterozygote Fittes (ii) Single Heterozygote Fittest (iii) Double Homozygote Fittest

With double heterozygote fittest (Figure 4i) and a low deterministic recombination fraction (e.g., *R* = 0.05), there are two complementary polymorphisms. However, once the recombination fraction surpasses a distinct *R* range threshold—dependent on the fitness matrix—the genotype distribution converges to both the corners and the central point [0.25, 0.25, 0.25, 0.25] depending on **x**_**0**_ (see Feldman and Liberman 1979).

With random *R*, a “convergence diagonal” is described by the parametric equations:

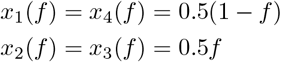

where *f* ∈ [0, 1].

Distinct transitions in convergence were observed as the range of random recombination fraction, *R*, expanded. Specifically:

1. For the initial increase in the range of *R*, the convergence regions grow across the parametric convergence diagonal (marked in dashed green).
2. When *R* increases beyond a particular range threshold, the genotypic distribution converges to the corners (see Figure 7i).
3. Upon exceeding the next range threshold, a central equilibrium becomes an attractor with a genotype distribution of [0.25, 0.25, 0.25, 0.25] (see Figure 7ii).

**Fig 5.**
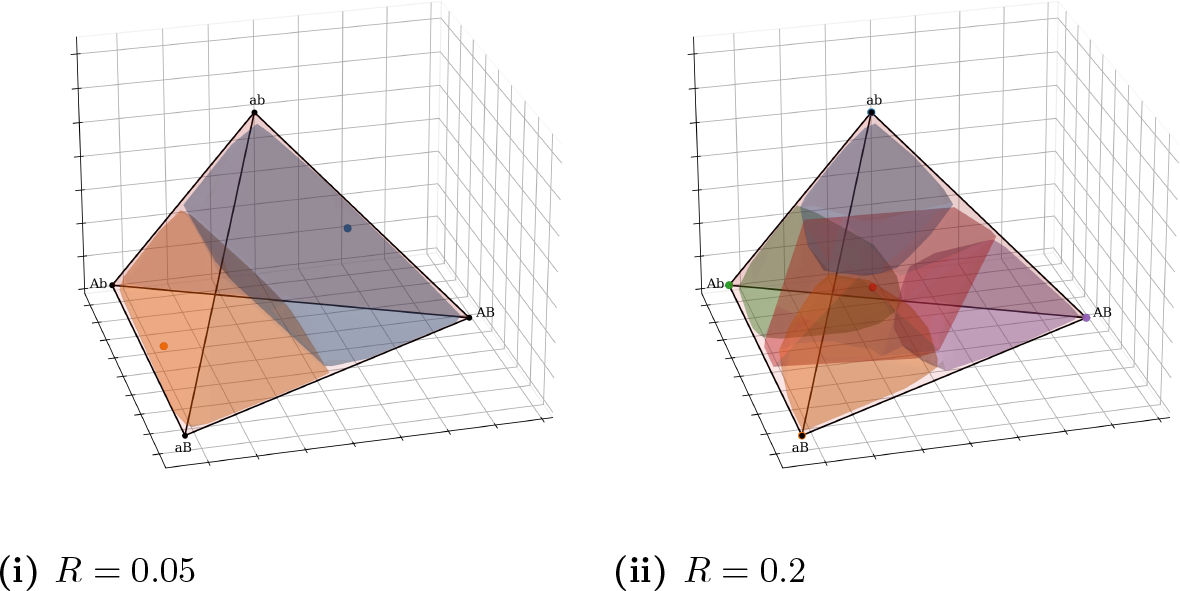
Convergence with Double Heterozygote Fittest and Deterministic *R*

**Fig 6.**
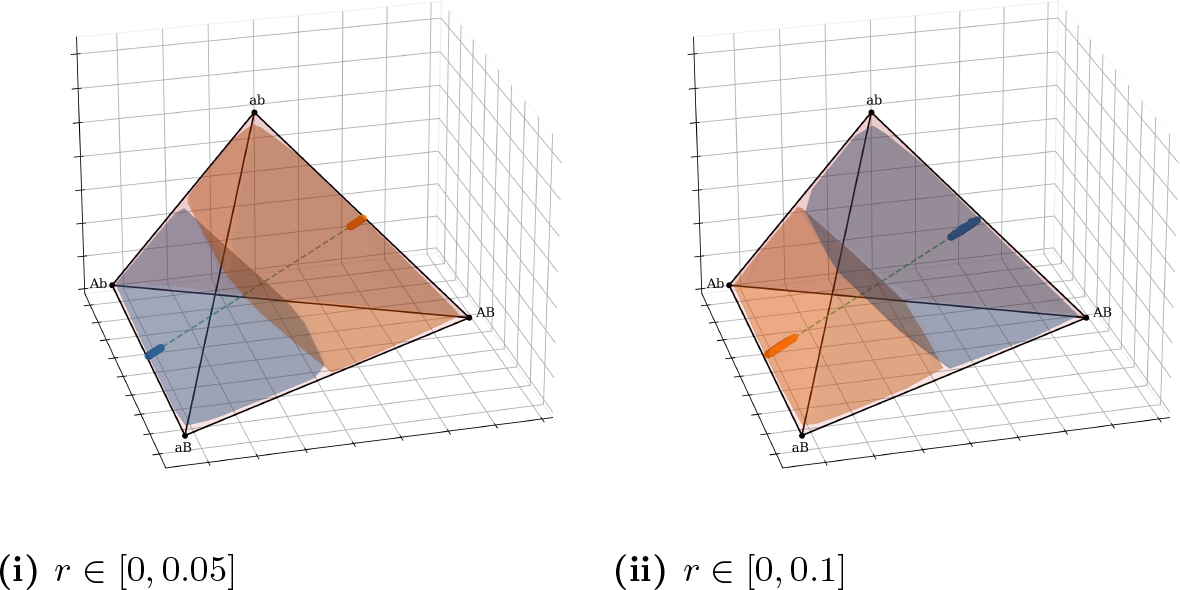
Convergent Region Inward Expansion Across Parametric Diagonal

**Fig 7.**
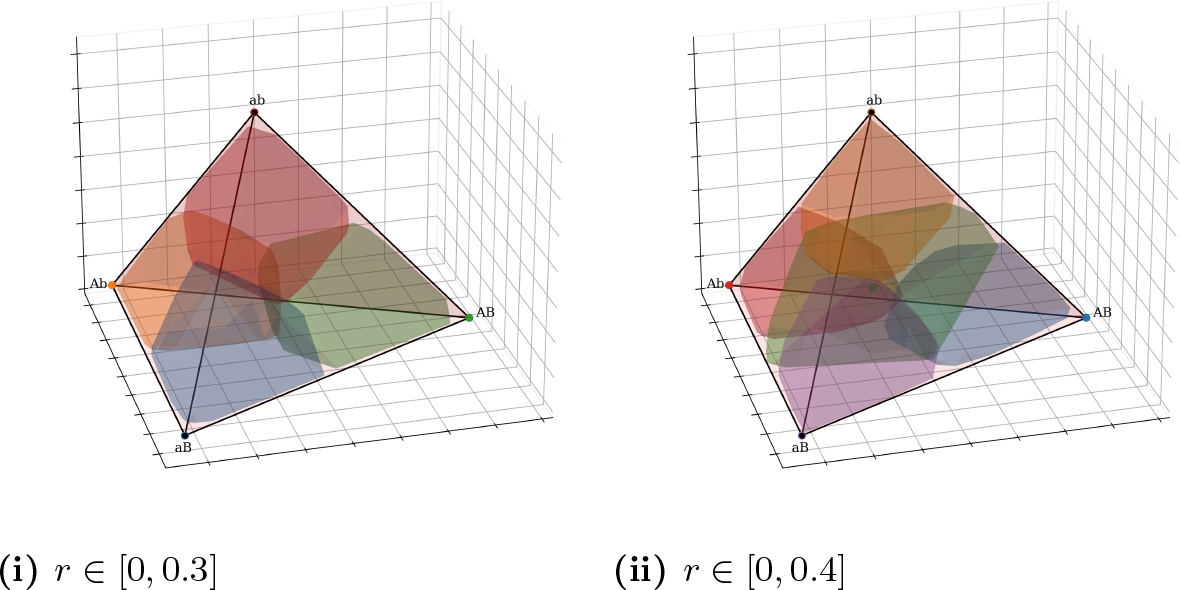
Convergence Progression for Expanding *R* Ranges

### 4.1 Single Heterozygote Fittest

In the deterministic model with single heterozygote fittest (Figure 4ii), the genotypic distribution approaches fixation to the corners. As the recombination fraction increases, the domain of **x**_**0**_ that converges to the corners aligns with the parametric diagonal (see Figure 8). Random recombination does not influence the genotypic outcomes.

**Fig 8.**
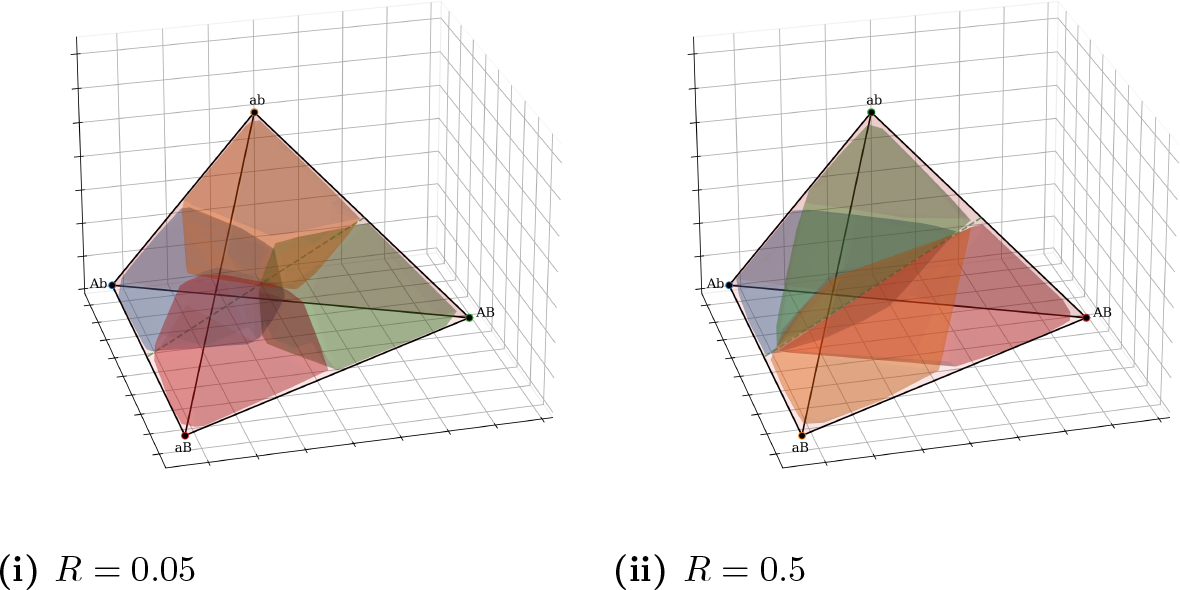
Convergence with Single Heterozygote Fittest Matrix and Deterministic *R*

### 4.2 Homozygote Fittest

With homozygote fittest (Figure 4iii), the deterministic, random, and no recombination scenarios converge to all non-homozygote edges.

**Fig 9.**
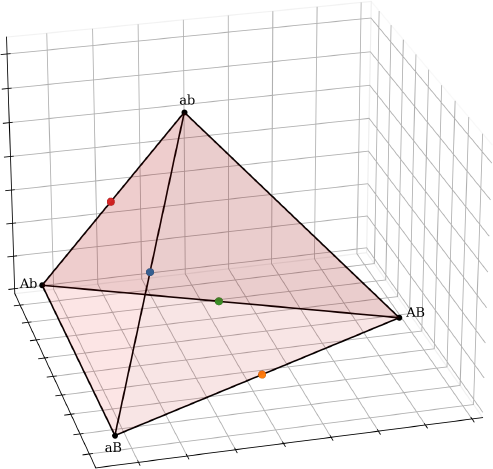
Homozygote Fittest Convergence

## 5 Convergence behavior under random fitness matrices

Dynamics in genetic systems have traditionally been understood within the context of deterministic fitness matrices. With fixed recombination, these matrices ensure that the trajectories of genotypic frequencies arrive at predictable outcomes. It would be natural to assume that introducing stochasticity to the fitness matrix — for example by resampling each element *w*_*ij*_ from a uniform distribution [0, 1] at every discrete time step — would disrupt these convergent trajectories.

With a random matrix without recombination, genotypes exhibit a low but consistent distribution throughout the tetrahedron’s interior. Recall from earlier that this spread-out distribution is attributed to a randomly occurring trend from the stochastic fitness elements, occasionally strong enough to drive the genotype toward the center. Genotypes exhibit a higher tendency towards the tetrahedral edges, a distribution caused by linkage disequilibrium. However, the highest concentration of genotypes is at the tetrahedral corners.

As the recombination fraction increases, the genotypes proximal to the *AB/ab* and *Ab/aB* edges experience a repulsion. Given that edges represent closely linked alleles, increased recombination disrupts these linkages, driving genotypes away. Random recombination does not influence genotypic outcomes; the dynamics behave analogously to those with a deterministic *R* equivalent to the mean of such a distribution.

**Fig 10.**
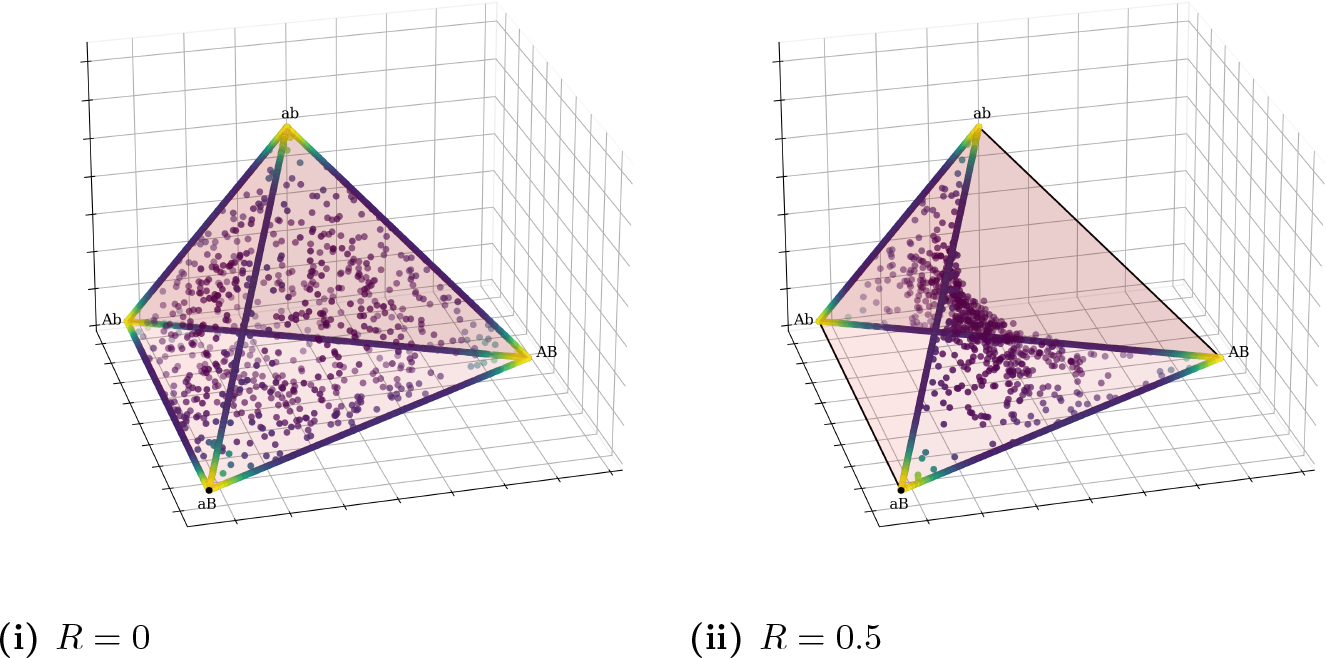
Convergent Distribution Density as *R* Increases

In the stochastic setting, the dynamics of the process can be modeled as a Markov chain on the space of non-degenerate frequency vectors

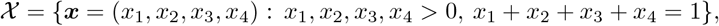

If the fitness values are i.i.d from the uniform distribution on (0, 1), one would expect that the trajectories of the process “cover” the full state space; this is in contrast with the deterministic model, where trajectories are fixed by the fitness matrix and the initial state. An immediate consequence is that fixed equilibria, i.e. frequency vectors which remain invariant over time, do not exist for the stochastic model. In fact, the only *global* equilibrium points (i.e. equilibria for *any* possible fitness matrix) in the *extended* state space (obtained by including the boundary of the tetrahedron) are the four corners; this follows from inequality (4). Therefore, should some equilibrium exist for the stochastic model, it should come as no surprise if that would imply these corners (which, importantly, do *not* belong to the interior 𝒳). Nevertheless, equilibria for Markov chains should be understood in a probabilistic sense and existence of such equilibria is the main concern of our analysis.

Formally, a Markov chain is a sequence of random variables {***X***_*n*_ : *n* ≥ 0} with a particular type of dependence. It is defined by a *transition probability* on the state space which gives the conditional probability distribution of the next state, given the current one. We will formally denote that by

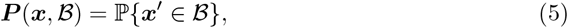

where ***x***^*′*^ is the random vector generated by (1) and *ℬ* is a (possibly open) subset of the state space 𝒳. The transition operator (5) together with the initial state ***X***_0_ determine the probabilistic behavior of the chain, just like the fitness matrix together with the initial state determine the trajectory of the deterministic process. We denote by ℙ _***x***_ the probability distribution of the chain started in ***X***_0_ = ***x***.

For any set *ℬ* ⊂ *𝒳* we define the *hitting time* of *ℬ* as the minimum random number of steps the process requires to first reach *ℬ*; in the formula,

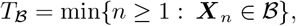

where the minimum is infinite if the chain never reaches set *ℬ*. If the initial state ***X***_0_ is a generic point in ℬ then *T* _*ℬ*_ may be also referred as *return time* to set ℬ. The set ℬ is called *accessible* from ***x*** if

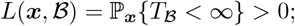

that is, if the chain started in ***x*** reaches set ℬ with positive probability. When ℬ is some open ball, which we shall denote by ℬ (***z***, *ϵ*) around ***z***, of radius *ϵ >* 0, we say that state ***z*** is accessible from ***x***. The Markov chain is said *irreducible* if for any pair of states ***x, z*** ∈ *𝒳* there exists some *ϵ >* 0 such that *L*(***x***, *ℬ* (***z***, *ϵ*)) *>* 0; in words, any state is accessible from any other state.

Irreducibility plays an important role in the analysis of a Markov chain. It ensures that, in principle, the chain will essentially visit any state in the state space, no matter what the initial state is. Indeed, this is the behavior we see in the generational limit in Figure 1.

To establish irreducibility of the Markov chain ***X*** we need to check under which circumstances there exist fitness values *w*_*ij*_ ∈ (0, 1) to satisfy (2), for some given values ***x, z*** = ***x***^*′*^∈ *𝒳* . Furthermore, note that solving the equation ***z*** = ***x***^*′*^ amounts to solving the system of equations

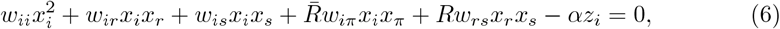

for *i* = 1, 2, 3, 4. There are 4 equations and 10 variables, all of them constrained to be in (0, 1). A general solution to this system can be found by assigning random values in (0, 1) to the variables *w*_*pq*_, with *p ≠q*, and then letting (*i* = 1, 2, 3, 4)

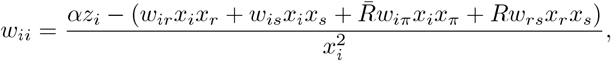

where we treat *α* ∈ (0, 1) as a parameter. Now we only need to impose that the calculated values *w*_*ii*_ meet the constraints *w*_*ii*_ ∈ (0, 1); this is because the last constraint, *α* ∈ (0, 1), will be then implicit since ***z*** ∈ *𝒳* entails

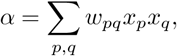

hence *α <* (*x*_1_ + *x*_2_ + *x*_3_ + *x*_4_)^2^ = 1, provided that ***x*** ∈ *𝒳* . The constraint *w*_*ii*_ *>* 0, for *i* = 1, 2, 3, 4, amounts to

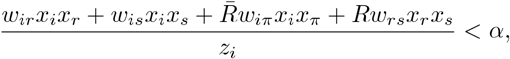

while *w*_*ii*_ *<* 1 is equivalent to

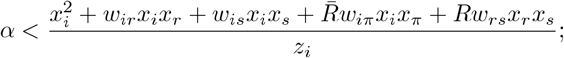

so, to make sure that the system has at least one solution, we require that there exist fitness values *w*_*pq*_ ∈ (0, 1), for *p ≠ q*, satisfying

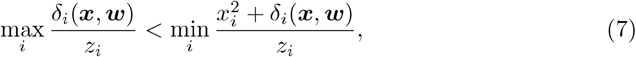

where we denote 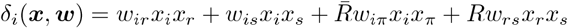. By a continuity argument, equation (7) gives sufficient conditions for state ***z*** being accessible, in *one* transition, from a state ***x***. To summarize our analysis, for any ***x, z*** ∈ *𝒳*, if there exist *w*_*pq*_ ∈ (0, 1), for *p ≠q*, satisfying (7), then

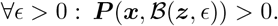

With this result in hand, we can go now further to establish irreducibility. More specifically, we will show first that:

I. the center ***a*** is accessible from any other state;
II. any state is accessible from a small neighborhood of the center ***a***.

For part (I) take ***z*** = ***a*** in (7) and, since *z*_*i*_ = 1*/*4, we need to show that

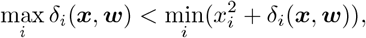

for some *w*_*pq*_ ∈ (0, 1). Taking 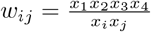 for *j* ∈ {*r, s*}, this reduces to

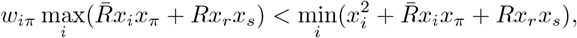

which obviously holds true for small enough *w*_*iπ*_.

By continuity of *δ*_*i*_ (in both ***x*** and ***w***), for part (II) it is enough to show that any state ***z*** is accessible from the center ***a***; then the conclusion will follow true, as well, for at least some small neighborhood around ***a***. Taking now ***x*** = ***a*** in (7), we need to find values *w*_*pq*_ ∈ (0, 1) such that

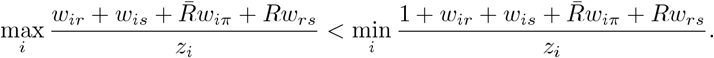

When *w*_*pq*_ = *ε*, for *p ≠ q*, for some *ε >* 0 (to be chosen later), this reduces to

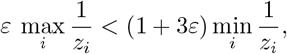

which is obviously true for *ε <* (min_*i*_ *z*_*i*_)*/*(max_*i*_ *z*_*i*_); this proves the claim.

Finally, we are able to prove irreducibility: take some arbitrary ***x, z*** ∈ *𝒳* and *ϵ >* 0. Then, by part (I) ***P*** (***x***, *𝒱*) *>* 0, for any neighborhood *𝒱* of ***a***, and by part (II) ***P*** (***v***, *ℬ* (***z***, *ϵ*)) *>* 0, for all ***v*** in some neighborhood *𝒱* of ***a***.

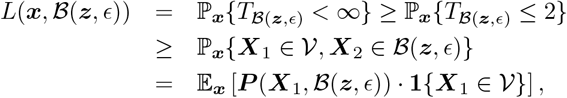

and the latter is strictly positive since it is the expectation of some positive random variable over a set of non-null probabilities. This proves that any state ***z*** may be reached from any ***x***_**0**_, with positive probability, in at most 2 steps, hence irreducibility follows true.

The high genotype concentration at the tetrahedral corners can be attributed to the behavior of genotype frequencies when approaching fixed points. To make this intuition precise, let us introduce the following terminology: let

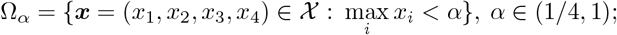

the sets Ω_*α*_, for *α <* 1, will be generically called *central regions*. The complement of the central region Ω_*α*_ can be partitioned as follows: ∁Ω_*α*_ = ∪_*i*=1:4_∆_*i,α*_, where ∆_*i,α*_ are disjoint (for *α >* 1*/*2), defined as ∆_*i,α*_ = {***x*** : *x*_*i*_ ≥ *α}*; these are *corner regions*, i.e. neighborhoods of the corresponding corners, and are getting small when *α* → 1. The sets Ω_*α*_ are increasing w.r.t. *α* ∈ (1*/*4, 1), so one can define the (set) limits inf_*α*_ Ω_*α*_ and sup*α* Ω_*α*_; the former is simply the *center* of the space ***a*** = (1*/*4, 1*/*4, 1*/*4, 1*/*4) and the latter is the standard (closed) simplex in R^4^ without the four corners. Thinking of central regions Ω_*α*_ as open, bounded neighborhoods of the center ***a***, we see that the four corners of the tetrahedron appear as the points that are not covered by any “bounded” region Ω_*α*_. The chain consistently interacts with the corner regions, culminating in a high concentration in these areas over time.

Proving the recurrence of the Markov chain extends beyond our Monte Carlo simulations and presents a fruitful avenue for future investigation. The stochastic equilibrium observed in Figure 1 is only possible in the limit of the chain is indeed recurrent. In this context, a stochastic equilibrium is understood as a probability distribution *μ* on *𝒳* which is left invariant by the transition operator – that is, if ***X***_0_ ∼ *μ* then ***X***_*n*_ ∼*μ* for any *n* ≥1. Nevertheless, for practical purposes, the generational span of our simulations should faithfully represent the dynamics of an actual population.

## 6 Discussion

Our investigation focused on the effect that random recombination rates and fitness matrices have on dynamics in the two-locus biallelic model. For deterministic trajectories, we identified a two-fold dynamic where genotype frequencies would move toward the center of the simplex before approaching stable polymorphisms.

Upon the introduction of randomly varying recombination, we identified regions of attraction *S*, to which ***x*** remains restricted with a high probability. Notably, the quantity and spatial distribution of these regions are contingent upon the fitness matrix and the range of the random recombination fraction. As the range of the random recombination fraction is increased, there are transitions in the location of the regions of attraction with both double and single heterozygote fittest cases from the edges across the parametric diagonal. In the homozygote fittest case, all recombination scenarios lead to fixation of the genotypes *AB* and *ab*. Further analysis with random fitness matrices demonstrated that genotype fixation acts potently as an attractor, drawing a high concentration of genotypes toward the tetrahedron’s vertices.

## 7 Acknowledgments

Some of the computing for this project was performed on the Sherlock cluster. We would like to thank Stanford University and the Stanford Research Computing Center for providing computational resources and support that contributed to these research results. Special thanks to Professor Marcus Feldman for his invaluable guidance and advice throughout this project.

## References

1. Fisher RA. The Genetical Theory of Natural Selection. Oxford University Press; 1930.

2. Wright. The Genetics of Quantitative Variability. London: Her Majesty’s Stationery Office. 1952;1(1):5–41.

3. Feldman, Karlin. Linkage and Selection: Two Locus Symmetric Viability Model. Theoretical Population Biology. 1970;1(1):39–71.

